# Effect of Aqueous Extract of Antiaris Africana Stem Bark on Phagocytic Activities of Human Neutrophil Ex-Vivo

**DOI:** 10.1101/2021.03.21.436291

**Authors:** Ibrahim O. Bello, Adebayo L. Adedeji

**Affiliations:** Nigerian Navy School of Health Sciences, Offa, Kwara State, Nigeria; Ladoke Akintola University of Technology, Ogbomoso, Oyo State, Nigeria

## Abstract

The immune system is one of the most complex biological systems in the body. During infection, the immune system is under attack by a large number of viruses, bacteria, fungi and parasites. Immune response firstly involves, recognition of the pathogen or foreign object and secondly, a reaction to eliminate it. Aqueous extract of *Antiaris africana* is used to study their immune modulator activity. This plant has its various parts used in folk medicines. The mechanism of action has not been fully elucidated. Therefore, this research work studies the effect of aqueous extract of *Antiaris Africana* stem bark on phagocytic activities of neutrophil isolated from apparently healthy individual using a non-subjective commercial colourimetric assay kit obtained from Cell-Biolab Inc., USA. The purity and viability of isolated neutrophils were >90% and >95% respectively. The extract enhances neutrophil phagocytosis at 1.0, 5.0, 10.0, 15.0, 20.0 and 25 μg/ml by 2.5%, 11.6%, 18.4%, 24.4%, 31.2% and 38.2% respectively, compared to the control (100%). Hence, it was observed that neutrophil phagocytosis increases with increased extract concentrations. It can be concluded from the study that enhancement of phagocytosis may be the possible mechanism of action of the plant as an immune modulator.

## INTRODUCTION

Phagocytosis is one of the most important arms of host defence, which helps to trigger host defence against invading pathogens. Phagocytosis is a biological process that consists of series of events, beginning with the binding and recognition of particles by cell surface receptors followed by the formation of actin-rich membrane extension and the particle (Larsson *et al.*, 2001).Since human body is constantly engaged in a bitter internal war against wide armies of microscopic enemies, phagocytosis by phagocytes prevent the body from pathogen infections (*Roitt et al.*, 1996). Phagocytosis is essential for a variety of biological events, including tissue remodelling and the continuous clearance of dying cells among others (Larsson *et al.*, 2001). The phagocytic process can be separated into several major stages such as chemotaxis, attachment of particles to the cell surface of phagocytes, ingestion and intracellular killing (*Sawyer et al.*, 1989). The phagocytic cells include neutrophils, eosinophils, and monocytes that recognise foreign substances and invading microorganisms.

Medicinal plants play important roles in various diseases and various plants have been shown to have use in pharmaceutical development. Traditional knowledge to solve health problems of man exists in all countries of the world (Rukangira, 2001). In most of the traditional medicine, the medicinal plants include the fresh or dried part, whole, chopped, powdered or an advanced form of the herb usually made via extraction by a solvent such as water, ethanol, or an organic solvent play a major role and constitute the backbone of traditional medicine (Mukherjee, 2002). The exploration of the chemical constituent of the plants and pharmaceutical screening may provide us the basis for the developing the lead for development of novel agents. Herbs have provided us some of the very important life-saving drugs used in modern medicine. Research have shown different plants used to solve immunodeficiency diseases. Also plants that enhance phagocytosis (Benencia *et al.*, 1994), inhibit phagocytosis (Atal *et al.*, 1995), and plants with no phagocytic effect (Simons *et al.*, 1990) has been reported.

The World Health Organisation has estimated that about 80% of the world’s population in developing countries use herbal remedies to treat various ailments (Akerele, 1994). This research study investigates the effect of aqueous extract of *Antiaris africana* on phagocytosis using neutrophil. *Antiaris africana* has its various parts used for numerous purposes in folk medicines (Banso and Mann, 2008). Research has been done using stem back extract of *Antiaris africana* on its antibacterial effect (Banso and Mann, 2008).

*Antiaris africana* is a plant found in various parts of Nigeria and West Africa. It is commonly called Ooro, Oriro or Ako Iroko in the South Western part of Nigeria; Farin Loko in the North; and Ojianwu in the South East. The plant is a large tree usually about 15 to 20 m high, but it can grow sometimes up to 40m, and has white latex and alternate dissymmetric leaves (Berg *et al.*, 1985; Berhaut, 1979), with heavy flat crown and blotchy grey and white bark. The flowers are small with a greenish white color that produces a red velvety fruits (Gill, 1992). It has a wide usage both in industry (timber *making) and traditional medicine. Various part of the plant such as leaves, stems and* barks are ethno-botanically used in the treatment of various diseases such as rheumatic and respiratory infection (Gill, 1992; Mann *et al.*, 2003), epilepsy, skin irritant, purgative, chest pain (Okogun *et al.*, 1976), syphilis (Berhaut, 1979), throat infection, leprosy, cancer (Kuete *et al.*, 2009), and nervous disorders in the northern part of Nigeria (Moronkola and Faruq, 2013).

## MATERIALS AND METHODS

### Materials

Spectrophotometer (HA-1600; West Sussex, BN15 8TN, UK), Rotary Evaporator (NYC, R-205D), *Antiaris africana* stem bark, deionised water, EDTA bottles, Centrifuge, weighing balance, needle and syringe, 15ml conical tube, micro-pipette, Neutrophil isolation kit, CytoSelect 96-Well Phagocytosis Assay kit obtained from Cell-Biolab Inc., USA.

## METHODS

### Plant Selection

The stem barks of *Antiaris africana* were collected in September, 2014 and was identified and authenticated at the Department of Forestry, Ekiti State Ministry of Environment, by the botanist – Mr. G.O Alo.

The bark was dried under shade, the dried bark was crushed into smaller pieces using mortar and pestle and was later grinded using a grinding machine.

### Aqueous Extraction of Antiaris africana

Fresh tree bark of *Antiaris africana* plant which was collected was washed to be free from sand and debris, cut into smaller pieces and dried under room temperature. The dried bark was crushed into smaller pieces using pestle and mortar and grinded using a grinding machine. The grinded dried bark was weighed using a weighing balance to be 500g and it was placed in a beaker of 2500ml and distilled water was added in ratio of 1:17 for 72 hours and stirred every 6 hours. The solution was further taken to the chemical laboratory for filtration process and extraction. The solution was sieved using muclin cloth and the residue was filtered using Buckner funnel. The filtered solution which is a clear but amber-yellow colour was poured into the rotary evaporator and it was evaporated at a temperature of 50°C and pressure ranges from 51 to 53mmHg for 24 hours. The evaporated solution further results in a dark soluble paste which was poured in a glass petri dish and placed in an oven at 50°C from where it turned into a powdered particle. Residue obtained after evaporation was weighed and percentage yield was determined using the formula; X/500g, where X=weight of residue after evaporation. The percentage yield was determined to be 38.3 %.

## PREPARATION OF NEUTROPHIL

5ml of whole blood was collected in an EDTA blood collection tube and was well labelled. The blood was transferred into a 50ml conical tube each. The blood collection tube was rinsed with 5ml of filtered cell-based assay buffer each. The rinsed solution was added to 50ml conical tube. 3.33ml of cell-based assay neutrophil isolation histopaque each was pipette to a different 50ml conical tube each. 10ml of diluted blood each was added on the top of cell-based assay neutrophil isolation histopaque. The samples were centrifuged at 500g for 20-30 minutes at 18-26°C. Aspiration was successfully done to remove the yellowish and clear top layers and leave the reddish pellet containing neutrophils and red blood cells in the tube. 5ml of the cell-based assay red blood cell lysis buffer into two different tubes. It was mixed using Vortex to ensure mixing of the cells with lysis buffer. It rocked in the rocker for 10-15 minutes to lyse the red blood cells. Centrifugation was done at 1,200 rpm for 10 minutes to pellet the neutrophils, and then the reddish supernatant was carefully aspirated. 1.66ml of RPMI containing 1% buffered saline was added to each tube and mix well. Centrifugation at 1,200 rpm for 5 minutes to pellet the neutrophil was done. The cells were suspended in 6.66ml of RPMI containing 1% Bovine Serum Albumin (BSA). It was mixed well to ensure sufficient separation of the cells. The cells are then ready to be seeded and were sufficient for two 96 well culture plates at a density of 1 x 10^5 – 5 x 10^5 cells/well.

## EVALUATION OF PURITY

Leishman staining was performed to confirm the neutrophil morphology and determined the percentage purity. Few drops of neutrophil suspension was spread on a glass slide, allowed to air dry and stained with Leishman stain for 1 minute. Subsequently, the smear was immersed with phosphate buffer solution for 15 minutes. The slide was rinsed off with tap water, dried and examined under the light microscope at X100 magnification. Thus, the differential percentage of neutrophils was determined.

## EVALUATION OF VIABILITY

Trypan blue dye exclusion viability assay was performed. Blood was diluted with the Trypan blue dye in a 1:19 ratio. Improved Neubauer counting chamber was charged with the solution, allowed to settle and examined under the light microscope using X10 magnification. Living cells exclude the dye whereas dead cells take up the dye. The blue is easily visible and the cells were counted. Thus, the percentage of living cells (viable cells) is expressed relative to the dead cells.

## ASSAY PROTOCOL: SUSPENSION PHAGOCYTES

Phagocytic cells were harvested and re-suspended in culture medium at 0.2 – 1.0 x 10^6 cells/ml. 10 μl was seeded in each cell of a 96-well plate. Phagocytes were treated with desired activators or inhibitors. 10ml of Zymosan suspension was added to each well. It was mixed well and immediately the plate was transferred to a cell culture incubator for 15 minutes – 12 hours. The cell culture was removed by centrifugation and gentle aspiration. 200μl of cold 1 x phosphate buffer saline (PBS) was added to each well. The PBS solution was removed by centrifugation and aspiration. 100μl of fixative solution was added to each well; it was incubated for 5 minutes at room temperature. The fixative solution was removed by centrifugation and gentle aspiration. It was washed twice with 1xPBS. 100μl of pre-diluted 1 x blocking solution was added to each well. The plate was incubated for 60 minutes at room temperature on an orbital shaker. The blocking solution was removed by centrifugation and gentle aspiration, then it was washed thrice with 1 x PBS. 100μl of pre-diluted 1x permeabilization solution was added to each well, and then incubated for 5 minutes at room temperature. The permeabilization solution was removed by centrifugation and gentle aspiration and then wash twice with 1x PBS. 100μl of pre-diluted 1x Detection reagent was added to each well. Incubation of the plate was done for 60 minutes at room temperature on an orbit shaker. The detection reagent solution was removed by centrifugation and gentle aspiration and then washed thrice with 1x PBS. 50μl of detection buffer was added to each well incubated the plate for 10 minutes at room temperature on an orbital. The reaction was started by adding 100 of substrate. Incubation for 5-20 minutes at 37°C was done. The reaction was stopped by adding 50μl of the stop solution mixed by placing the plate on an orbital plate shaker for 30 minutes. The absorbance of each well was read at 405nm.

## RESULT

### Plant Selection and Collection

After the selection of the plant, it was identified and authenticated by the Botanist, Mr. G. O Alo of Ekiti State Ministry of Environment, Ado Ekiti, Ekiti State.

### Purity

The morphological observation revealed the neutrophils with intact cell membrane, having 2-5 lobes of nucleus which stained dark blue, joined together with fine strands of chromatins and the cytoplasm stained clear light pink consisting of granules. Based on Leishman staining, >90% of the cell isolated were neutrophils.

### Viability

Trypan blue white cell viability analysis showed that >95% of the isolated neutrophils were viable. That is, after the isolation, viable (unstained) cells were determined to be about 95% of total cells.

### Phagocytosis Assay

Aqueous extract of *Antiaris africana* inhibit the phagocytic activity of neutrophil ex-vivo at different concentrations of 1.0, 5.0, 10.0, 15.0, 20.0 and 25.0 μg/ml.

Table 1 shows the mean ± standard deviation of the calculated absorbance gotten from three absorbance values, the volume of extract added and % change in phagocytic activity by the treatment with the extract at different concentrations. 100% phagocytic activity was observed at 0 μg/ml concentration which served as the control. Increase in phagocytic activity was observed at every increase in extract concentration. Highest increase was observed at 25μg/ml (7.4%) while the least increase was observed at 1μg/ml (1.4%). The increase in phagocytosis at all extract concentration were statistically significant (p<0.05) when compared against the control.

**Table 1:** Mean ± SD of Absorbance and % Change in Phagocytic Activity.

| CONC. OF EXTRACT | VOL. OF EXTRACT | MEAN ± SD OF ABSORBANCE | % INCREASE IN PHAGOCYTIC ACTIVITY | p-VALUE |
| --- | --- | --- | --- | --- |
| 0µg/ml | Cell only | 1.644 ± 0.012 | 100% | ----- |
| 1µg/ml | 0.5µL | 1.658 ± 0.020 | 1.4% | 0.001* |
| 5µg/ml | 2.5µL | 1.669 ± 0.029 | 2.5% | 0.000 * |
| 10µg/ml | 5.0µL | 1.682 ± 0.041 | 3.8% | 0.003* |
| 15µg/ml | 7.5µL | 1.693 ± 0.055 | 4.9% | 0.000 * |
| 20µg/ml | 10.0µL | 1.705 ± 0.068 | 6.1% | 0.000 * |
| 25µg/ml | 12.5µL | 1.718 ± 0.081 | 7.4% | 0.000 * |

\*p-value <0.05 is considered significant.

## DISCUSSION AND CONCLUSION

### Discussion

The immune system comprises of two mechanisms, primary innate response and; secondary adaptive immune response. Though adaptive response is highly specific to the antigens, non-specific innate response play an important role in the initial stages of defence, and one of its main attributes is phagocytosis. Phagocytic cells engulf and destroy foreign substances. Cells which possess such activities are neutrophils, eosinophils, macrophages and monocytes.

Phagocytosis by phagocytes is essential for a variety of biological events, including tissue remodelling and the continuous clearance of dying cells. Phagocytosis comprises a series of events, starting with the binding and recognition of particles by cell surface receptors till it ends in pathogen inside the phagolysosome formed are destroyed.

Neutrophils are short-lived and highly active cells where the isolation of neutrophils requires careful steps to yield a good amount of cells within a shorter period of time. To characterize the specific functions of neutrophils, a high purity, fast and reliable method of separating them from other blood cells is desirable for in-vitro studies.

In this study, 90% of cell purity was achieved. Possible reasons for not been able to achieve a 100% purity might be because of the little differences in the relative mass densities of blood cells, as the isolation method used in this study is density gradient method. Neutrophils and lymphocytes have a close mass density values, thus this increases the possibility of lymphocyte contamination.

Also, vigorous centrifugation of blood pushes lots of lymphocytes to the bottom of tube and contaminate purity of isolated neutrophil cells. Long duration of isolation protocol is another factor that could increase the contamination and decreasing the viability of isolated cells (Rezpour and Jaafar, 2009). In this study, >95% viability of isolated neutrophil cells was achieved. Neutrophils are short-lived, thus long isolation protocol and delay in testing process could cause decrease in the viability of isolated cells (Rezpour and Jaafar, 2009). In this study however, the isolated neutrophil cells were tested immediately and there were no delay during the testing process. The reason for not been able to achieve the 100% viability might have arose from the fact that some of the cells might have been close to the end of their life span.

Many plant extracts and medicinal plants have been shown to have potent phagocytic activity (Bhanu *et al.*, 2012).

The study demonstrated that aqueous extract of *Antiaris africana* enhances neutrophil phagocytic activity ex-vivo. In this study, aqueous extract of *Antiaris africana* showed increase in phagocytic activities of neutrophil at all concentrations compared to the control (100%) in a concentration-dependent manner.

Similar results have been reported from other plants and several researchers have shown some plant extract with enhancement effect on phagocytosis (Benencia *et al.*, 1994).

This study supports previous work done on the plant by Kuete *et al.* (2009) which showed that the component compounds of *Antiaris africana* plant extract have anti-tumour and antioxidant activities.

However, some researchers using other plant extracts have reported no phagocytic activity (Simons *et al.*, 1990).

### Conclusion

This study shows that aqueous extract of *Antiaris africana* enhances phagocytosis in neutrophil. We therefore conclude, based on the results obtained from this research work, that enhancement of phagocytosis may be the possible mechanism of action of the plant.

